# Single-cell transcriptomics reveals multiple neuronal cell types in human midbrain-specific organoids

**DOI:** 10.1101/589598

**Authors:** Lisa M. Smits, Stefano Magni, Kamil Grzyb, Paul MA. Antony, Rejko Krüger, Alexander Skupin, Silvia Bolognin, Jens C. Schwamborn

**Author notes:** **Corresponding author** Prof. Dr. Jens C. Schwamborn.

## Abstract

Human stem cell-derived organoids have great potential for modelling physiological and pathological processes. They recapitulate *in vitro* the organisation and function of a respective organ or part of an organ. Human midbrain organoids (hMOs) have been described to contain midbrain-specific dopaminergic neurons that release the neurotransmitter dopamine. However, the human midbrain contains also additional neuronal cell types, which are functionally interacting with each other. Here, we analysed hMOs at high-resolution by means of single-cell RNA-sequencing (scRNA-seq), imaging and electrophysiology to unravel cell heterogeneity. Our findings demonstrate that hMOs show essential neuronal functional properties as spontaneous electrophysiological activity of different neuronal subtypes, including dopaminergic, GABAergic, and glutamatergic neurons. Recapitulating these *in vivo* features makes hMOs an excellent tool for *in vitro* disease phenotyping and drug discovery.

## Introduction

Current *in vitro* approaches to model physiology and pathology of human neurons are mainly based on pure cultures of neurons grown under 2D conditions. It has been shown that the differentiation potential of human induced pluripotent stem cells (iPSCs) provides a unique source of different neural cell types (Takahashi and Yamanaka, 2006). Until now, many protocols for generating iPSC-derived neural cultures have been described. The resulting cell culture monolayers have been proven as useful tools to study disease mechanisms and to identify potential neuroprotective compounds (Nguyen et al., 2011; Cooper et al., 2012; Sánchez-Danés et al., 2012; Reinhardt et al., 2013b; Ryan et al., 2013). However, these culture conditions does not recapitulate several characteristics, which are relevant to the human brain, like cyto-architecture or complex cell-cell interactions. This may result in inaccurate modelling of the human brain (patho-)physiology with the consequence that candidate compounds might prove efficacy in 2D *in vitro* studies but are ineffective in clinical trials or vice versa (Abe-Fukasawa et al., 2018). The recent establishment of new 3D neuronal cell culture models has contributed to mimic key aspects of human brain development (Lancaster et al., 2013; Tieng et al., 2014; Muguruma et al., 2015; Jo et al., 2016; Qian et al., 2016; Monzel et al., 2017). Studies using human cerebral brain organoids have shown the acquisition of neuronal maturity and network activity (Quadrato et al., 2017; Matsui et al., 2018). Their complex, multicellular architecture enables the study of neuronal diseases and has already led to novel insights on e.g. Zika virus-induced microcephaly (Ming et al., 2016; Qian et al., 2017). Besides this unique *in vitro* disease modelling potential, human brain organoids provide a platform for advanced drug screening (Kelava and Lancaster, 2016; Di Lullo and Kriegstein, 2017). In this study, we focused on a detailed characterisation of the different neuronal subtypes in human midbrain-specific organoids (hMOs). With single-cell transcriptome analysis, we examined the presence of different neuronal subtypes, and subsequently studied the effect of chemical compounds on the electrophysiological activity of the neuronal network. Our findings demonstrate that hMOs contain, beside dopaminergic neurons, other neuronal subtypes including GABAergic, glutamatergic, and serotonergic neurons. hMOs showed essential neuronal functional properties during the course of differentiation, like synapse formation and spontaneous electrophysiological activity. These features indicate that hMOs recapitulate specific characteristics of functional human midbrain tissue, thus making them a valuable resource for *in vitro* disease modelling and drug discovery.

## Material and Methods

### Ethics Statement

Written informed consent was obtained from all individuals who donated samples to this study and the here conduced work was approved by the responsible ethics commissions. The cell lines used in this study are summarised in Table 1.

**Table 1:**
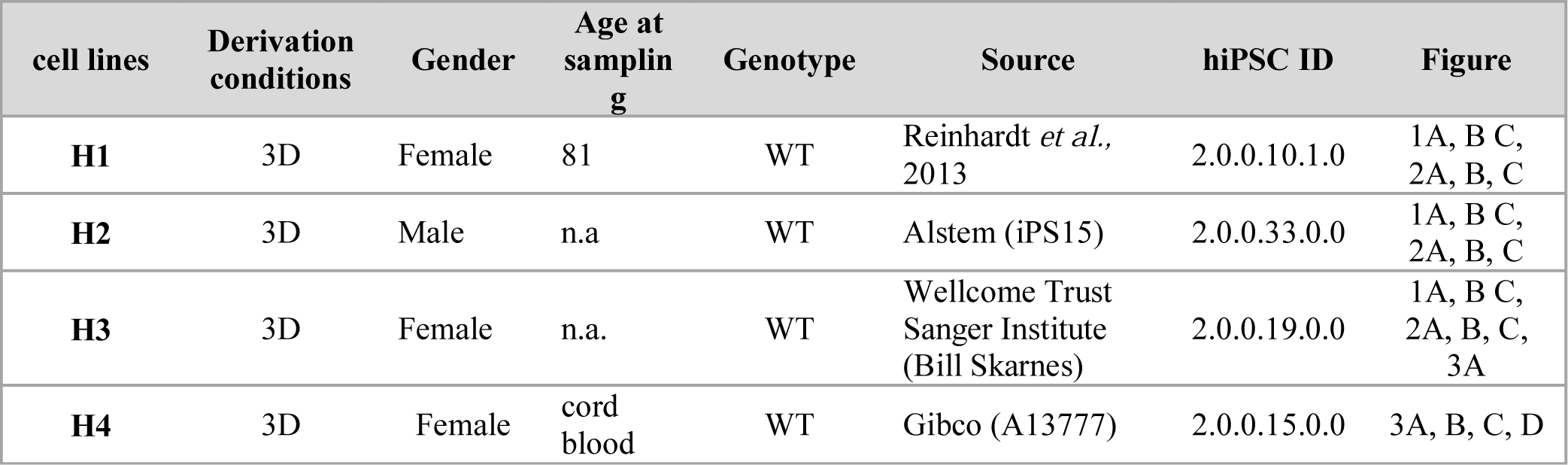
Cell lines used in this study to generate mfNPCs and midbrain-specific organoids. Human mfNPCs were derived under 2D conditions from human iPSCs of different origin. hMOs were generated as described in the experimental procedures section.

### Pluripotent Stem Cell culture

hiPSC lines were provided by Bill Skarnes, Wellcome Trust Sanger Institute (iPSC Bill), Alstem (iPS15, derived from human peripheral blood mononuclear cells, episomal reprogrammed) or previously described in Reinhard *et alia* (Reinhardt et al., 2013b). The cells were cultured on Matrigel-coated (Corning, hESC-qualified matrix) plates, maintained in Essential 8 medium (Thermo Fisher Scientific) and cultured with and split 1:6 to 1:8 every four to five days using Accutase (Sigma). 10 µM ROCK inhibitor (Y-27632, Abcam) was added to the media for 24 h following splitting.

### Derivation of midbrain floorplate neural progenitor cells

The derivation and maintenance of midbrain floorplate neural progenitor cells (mfNPCs), has been described previously (Smits et al., 2019).

In brief, embryoid bodies (EBs) were formed with 2,000 iPSCs each, using AggreWell 400 (Stemcell Technologies). The cells were cultured in Knockout DMEM (Invitrogen) with 20 % Knockout Serum Replacement (Invitrogen), 100 µM beta-mercaptoethanol (Gibco), 1 % nonessential amino acids (NEAA, Invitrogen), 1 % penicillin/streptomycin/glutamine (Invitrogen), freshly supplemented with 10 µM SB-431542 (SB, Ascent Scientific), 250 nM LDN-193189 (LDN, Sigma), 3 µM CHIR99021 (CHIR, Axon Medchem), 0.5 µM SAG (Merck), and 5 µM ROCK inhibitor (Sigma). After 24 h, EBs were transferred to a non-treated tissue culture plate (Corning). On day two, medium was replaced with N2B27 medium consists of DMEM-F12 (Invitrogen)/Neurobasal (Invitrogen) 50:50 with 1:200 N2 supplement (Invitrogen), 1:100 B27 supplement lacking vitamin A (Invitrogen) with 1 % penicillin/streptomycin/glutamine, supplemented with 10 µM SB, 250 nM LDN, 3 µM CHIR, 0.5 µM SAG. On day four and six, medium was exchanged with the same but including 200 µM ascorbic acid (AA, Sigma). On day eight, EBs with neuroepithelial outgrowth were triturated into smaller pieces and diluted in a 1:10 ratio. For following passages, 1x TrypLE Select Enzyme (Gibco)/0.5mM EDTA (Invitrogen) in 1x PBS was used and 10,000 to 20,000 cells per 96-well ultra-low attachment plate (round bottom, Corning) were seeded. The cells were always kept under 3D culture conditions and from passage 1 on cultured in N2B27 medium freshly supplemented with 2.5 µM SB, 100 nM LDN, 3 µM CHIR, 200 µM AA, and 0.5 µM SAG. After every cell split the ultra-low attachment plate was centrifuged for 3 min at 200 xg to assure the aggregation of single cells at the bottom of the well. Additionally, 5 µM ROCK inhibitor was added. The cells were split every 7 to 14 days and the medium was changed every third day. After four to five passages, mfNPCs were used as a starting population for hMOs.

### Generation of midbrain-specific organoids

To start the generation of hMOs, 3,000 cells per well were seeded to an ultra-low attachment 96-well round bottom plate, centrifuged for 3 min at 200 xg and kept under maintenance conditions for seven days. LDN and SB were withdrawn of mfNPC expansion medium and after three additional days, the concentration of CHIR was reduced to 0.7 µM. On day nine of differentiation, medium was changed to neuronal maturation N2B27 medium including 10 ng/ml BDNF (Peprotech), 10 ng/ml GDNF (Peprotech), 200 µM AA (Sigma), 500 µM dbcAMP (Sigma), 1 ng/ml TGF-β3 (Peprotech), 2.5 ng/ml ActivinA (Life Technologies) and 10 µM DAPT (Cayman). The organoids were kept under static culture conditions with media changes every third day for 35 or 70 days. Detailed information about the generation of hMOs has been published recently (Smits et al., 2019).

### Immunofluorescence

hMOs were fixed with 4 % PFA overnight at 4 °C and washed 3x with PBS for 15 min. After treatment, they were embedded in 3-4 % low-melting point agarose in PBS. The solid agarose block was sectioned with a vibratome (Leica VT1000s) into 50 µm sections. The sections were blocked on a shaker with 0.5 % Triton X-100, 0.1 % sodium azide, 0.1 % sodium citrate, 2 % BSA and 5 % normal goat or donkey serum in PBS for 90 min at RT. Primary antibodies were diluted in the same solution but with only 0.1 % Triton X-100 and were applied for 48 h at 4 °C.

After incubation with the primary antibodies (Table 2), sections were washed 3x with PBS and subsequently blocked for 30 min at RT on a shaker. Then sections were incubated with the secondary antibodies in 0.05 % Tween-20 in PBS for two hours at RT and washed with 0.05 % Tween-20 in PBS and Milli-Q water before they were mounted in Fluoromount-G mounting medium (Southern Biotech).

**Table 2:**
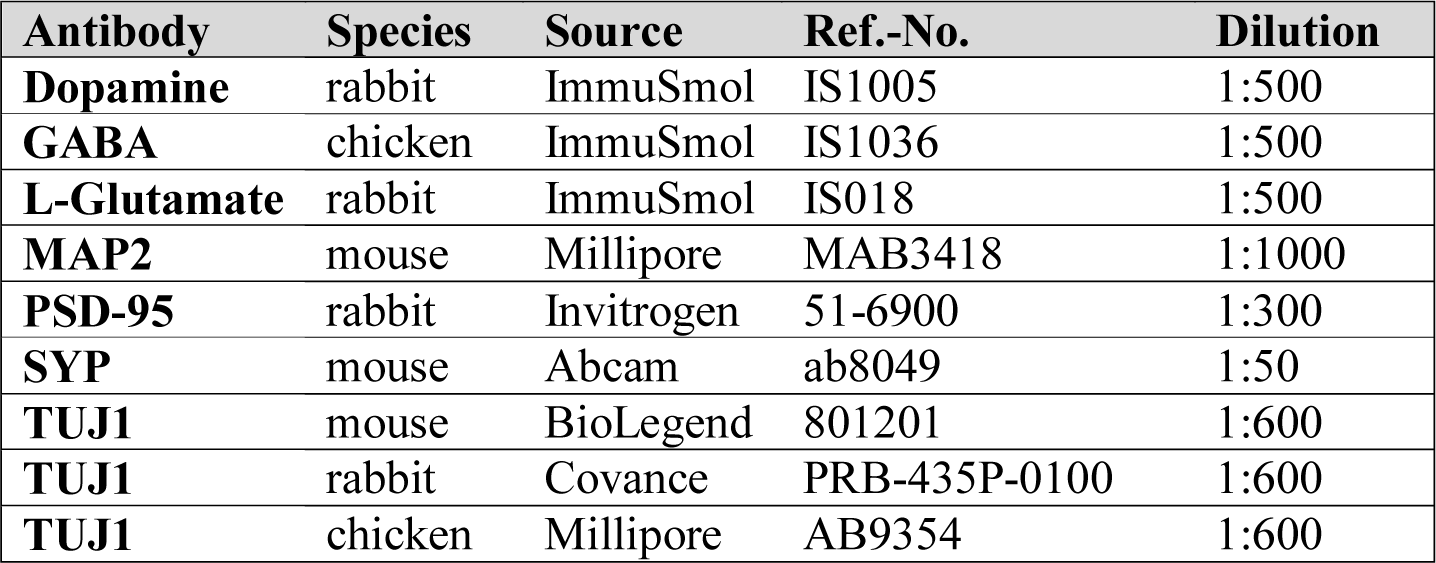
Antibodies used in this study.

STAINperfect Immunostaining Kit (ImmuSmol) was used according to manufacturer’s protocol to detect dopamine, serotonin, GABA and L-glutamine. Nuclei were counterstained with Hoechst 33342 (Invitrogen).

For qualitative analysis, three randomly selected fields per organoid section were acquired with a confocal laser scanning microscope (Zeiss LSM 710) and images were further processed with OMERO Software. Three-dimensional surface reconstructions of confocal z-stacks were created using Imaris software (Bitplane).

### Quantitative Image analysis

Immunofluorescence 3D images of hMOs were analysed in Matlab (Version 2017b, Mathworks). The in-house developed image analysis algorithms automate the segmentation of nuclei, astrocytes and neurons with structure-specific feature extraction. The image preprocessing for the segmentation of nuclei was computed by convolving the raw Hoechst channel with a gaussian filter. By selecting a pixel threshold to identify apoptotic cells, a pyknotic nuclei mask was identified and subtracted from the nuclei mask.

For the segmentation of neurons, a median filter was applied to the raw TUJ1 channels. The expression levels were expressed in two ways: i) positive pixel of the marker, normalised by the pixel count of Hoechst; ii) cells positive for a marker expressed as a percentage of the total number of cells. In this latter case, the nuclei were segmented and a watershed function was applied. Considering the high cell density of the specimens, steps to ensure high quality in the segmentation process were implemented and structure with a size higher than 10,000 pixels were removed (this indicated incorrected segmentation e.g. clumps). In the nuclei successfully segmented as a single element, a perinuclear zone was identified. In case the marker of interest was positive in at least 1 % of the perinuclear area, the corresponding cell was considered as positive.

### Single-cell RNA-sequencing using Droplet-Sequencing (Drop-Seq)

scRNA-seq data were generated using the Droplet-Sequencing (Drop-Seq) technique (Macosko et al., 2015) as described previously (Walter, 2019). In this work, we performed scRNA-seq of hMOs derived from hiPSC line H4 (see Table 1). For each time point, 35 d and 70 d after dopaminergic differentiation, we pooled and analysed 30 hMOs each.

### Pre-processing of the digital expression matrices from scRNA-seq

The result of the Drop-Seq scRNA-seq pipeline and subsequent bioinformatics processing is a digital expression matrix (DEM) representing the number of mRNA molecules captured per gene per droplet. Here, we obtained two DEMs, one corresponding to 35 d hMOs and the other to 70 d hMOs. After quality cut based on knee plots, we retained for each sample 500 cells with the highest number of total transcripts measured and performed normalisation of the DEM separately. Finally, the two DEMs were merged for the comparison analysis of the two time points based on 24,976 expressed genes in 1,000 cells. The data was analysed by our customized Python analysis pipeline (version 3.6.0, with anaconda version 4.3.1) including dimensionality reduction by t-distributed stochastic neighbourhood embedding (t-SNE) (van der Maarten and Hinton 2008) and differentially gene expression analysis.

### Analysis of DEGs from scRNA-seq data

To determine, which and how many genes were differentially expressed between 35 d and 70 d hMOs, we applied one-way ANOVA test, a one-way ANOVA test on ranks (Kruskal-Wallis test), and a Mutual Information based test. The minimum p-value obtained for each gene across these three tests was retained and statistical significance was set to p < 0.01 after Bonferroni correction for differentially expressed genes (DEG).

### Cumulative gene expressions from scRNA-seq data

From literature, we extracted cell-type specific gene lists (Table 3) for stem cells, neurons, and neuronal subtypes (dopaminergic, glutamatergic, GABAergic and serotonergic neurons) (Reinhardt et al., 2013a; La Manno et al., 2016; Cho et al., 2017). Note, that not all genes listed therein have been measured in our dataset, these were highlighted in Table 3.

**Table 3:**
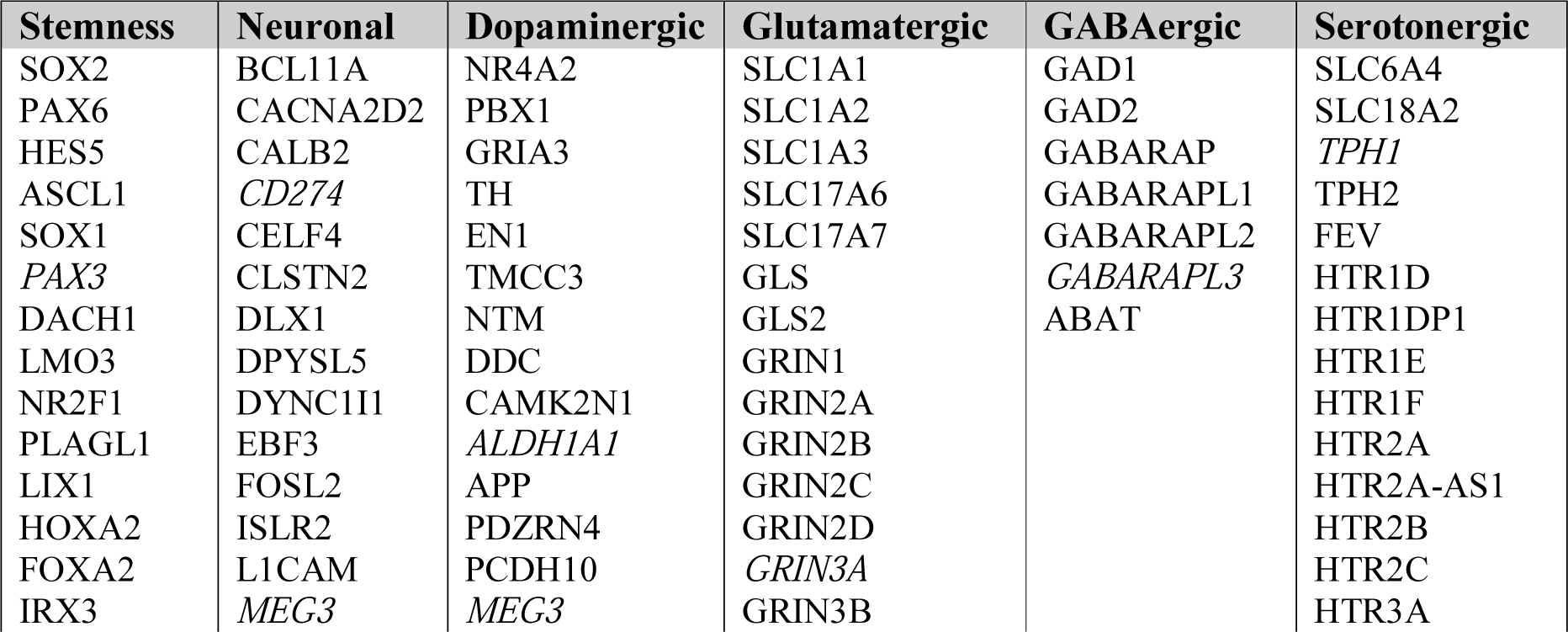
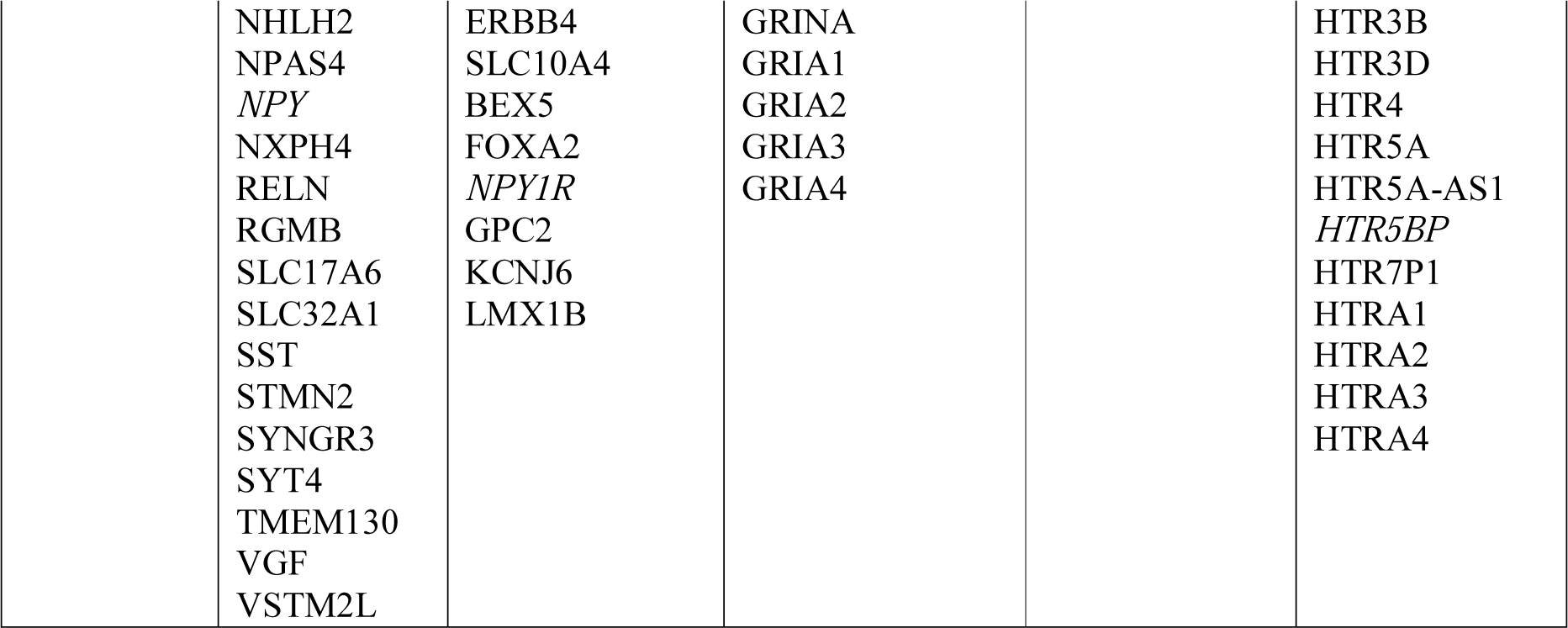
Gene lists used in this study. Genes that were not detected in the transcriptome are emphasised in italics.

For each list we defined a score, which we refer to as cumulative gene expression, computed as the sum of the corresponding genes from normalized DEM for each cell. Since the expression levels were measured at single cell level, we can consider the cells distributions across the cumulative genes expression scores (Figure 1D and Figure 2A). These histograms exhibit the cumulative gene expression scores normalised to their maxima on the horizontal axis. Thus, on the horizontal axis, a value of 1 corresponds to the maximal cumulative gene expression for one list of genes, while 0 corresponds to no expression of any genes from that list. The vertical axis exhibits the number of cells falling into the corresponding bin of the histogram. In each subpanel the distributions for day 35 and for day 70 are shown. Population differences were assessed by Z-test of the means with Bonferroni correction.

**Figure 1:**
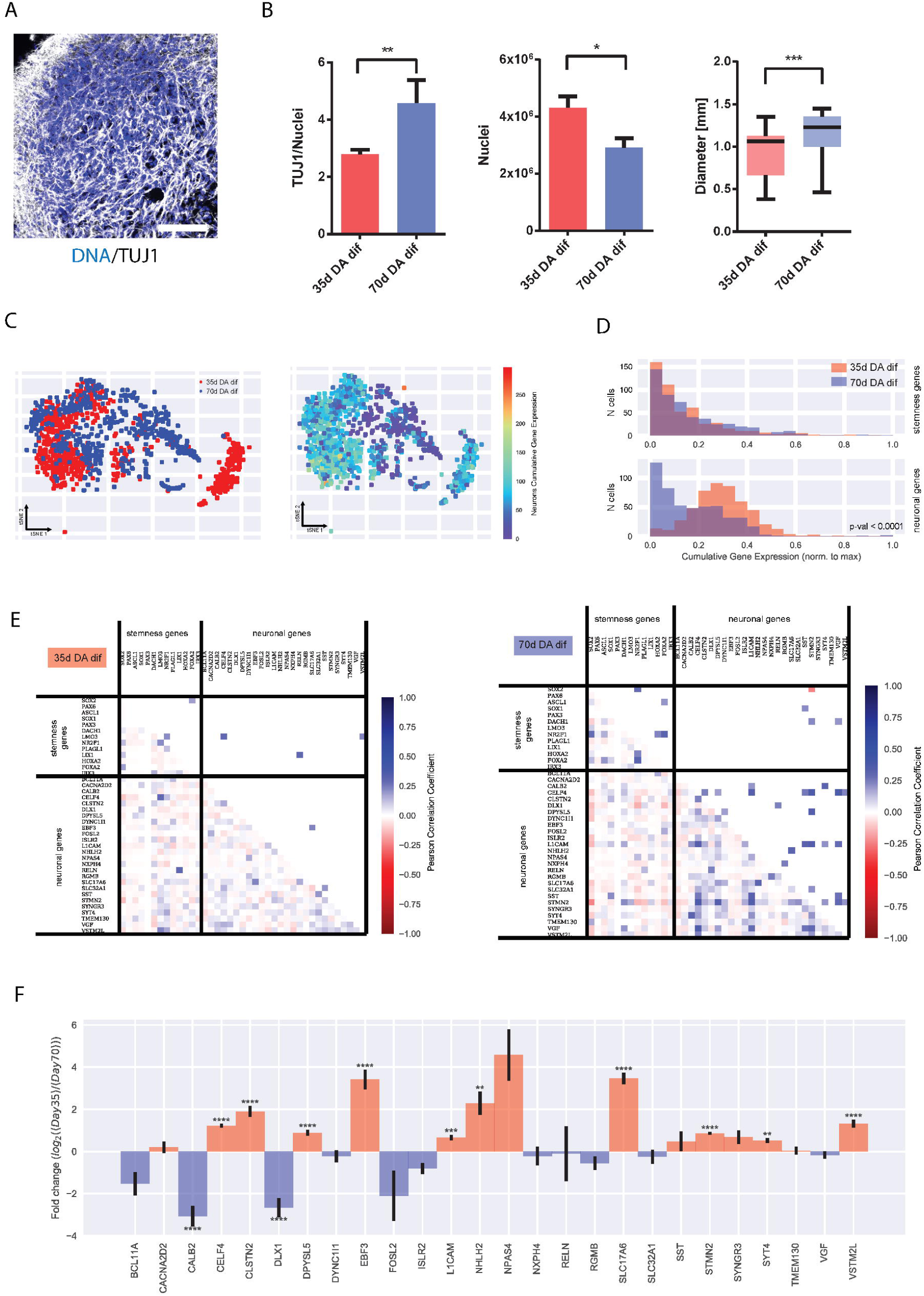
Identification of neuronal population in midbrain-specific organoids. (A) Immunohistological staining of TUJ1 expressing neurons in 35 d organoid sections (50 μm thickness, scale bar 100 µm). (B) The ratio of TUJ1 positive pixels normalised against Hoechst (35 d n=59, 70 d n=48). Quantification of Hoechst positive pixel (35 d n=22, 70 d n=29). Average size of four different organoid lines. Whiskers present minimum and maximum (35 d n=21, 70 d n=44). Data presented as mean ±SEM. (C) Dimensionality reduction of the scRNA-seq data by tSNE underlies differences in gene expression between the samples at day 35 and day 70. Each dot corresponds to a cell. In the left panel, colours are used to indicate cells from the two time points. In the right panel, the colour scale indicates the score (cumulative gene expression) corresponding to neuron-specific genes for each cell. (D) Distributions (histograms) of cells across the cumulative gene expression scores, obtained from lists of genes specific for precursor cells (stemness genes) or neurons (stemness genes). (E) Gene-gene correlation matrices, for genes at day 35 on the left, and day 70 on the right. (F) Log2 fold-changes between day 35 and day 70 in gene expression for individual genes corresponding to the lists for neuron-specific genes.

**Figure 2:**
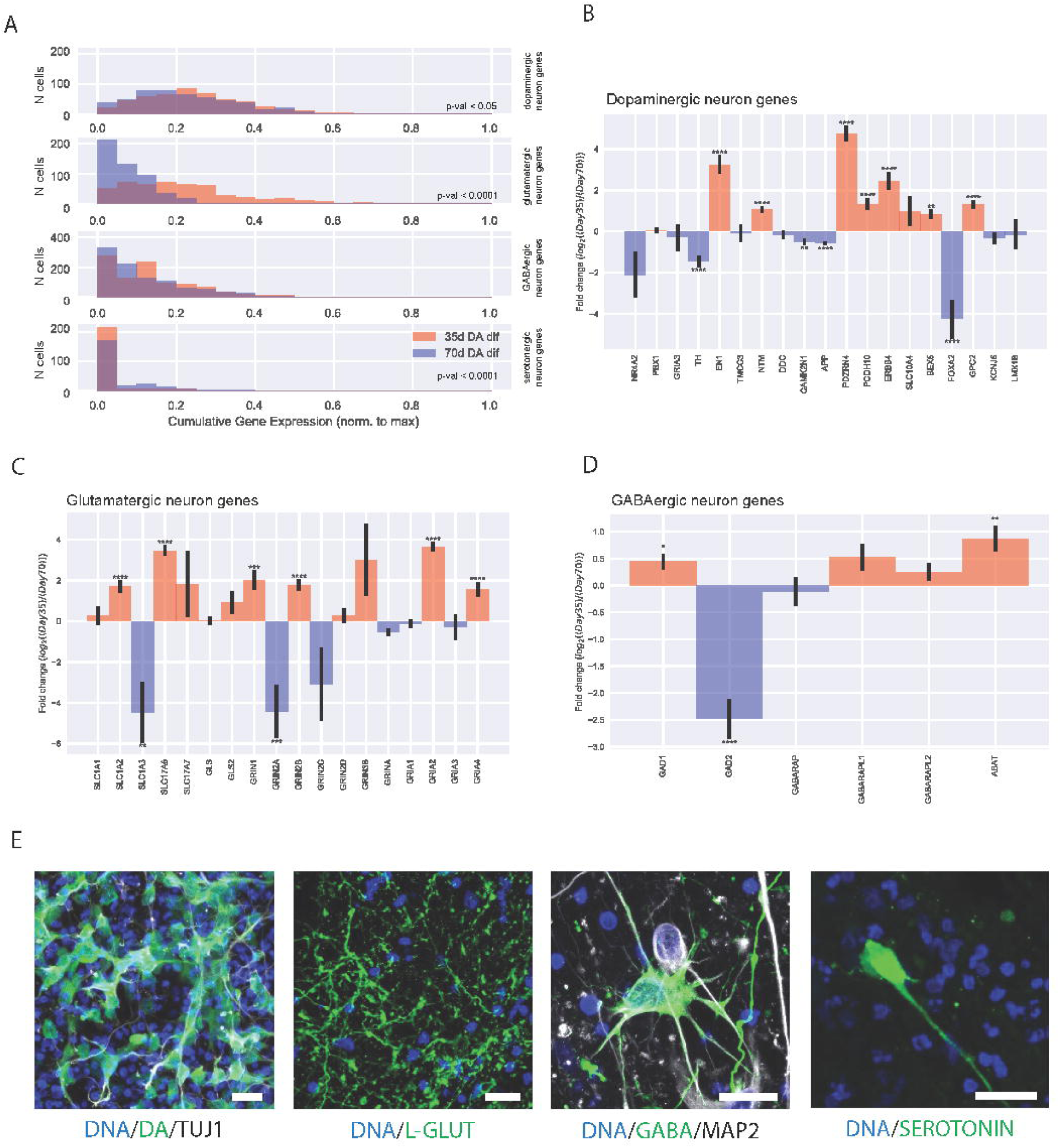
Neuronal subtypes in midbrain-specific organoids. (A) Distributions (histograms) of cells across the cumulative gene expression scores, obtained from the lists of genes specific for the main neuronal subtypes present in the organoids, namely dopaminergic, glutamatergic, GABAergic, and serotonergic neurons. (B-D) Log2 fold-changes between day 35 and day 70 in gene expression for individual genes corresponding to the lists of genes typical of the neuronal subtypes: (B) dopaminergic neurons, (C) glutamatergic neurons, and (D) GABAergic neurons. (E) Immunohistological staining of 70 d organoid sections (50 μm thickness). Detection of the neurotransmitters dopamine, L-glutamine, GABA, and serotonin. Scale bar is 20 μm.

### Gene-gene correlations from scRNA-seq data

From the scRNA-seq data we also computed gene-gene Pearson correlation coefficients for stemness- and neuron-specific genes. Analysis was performed independently for the two samples (35d DA dif and 70d DA dif) resulting in two correlation matrices (Figure 1E).

In the lower triangular matrix all correlation values are shown, whereas the upper triangular matrix only statistical significant correlations (p-value < 0.05 after Bonferroni correction). For visual clarity, diagonal elements and not detected genes were excluded.

### Fold changes of gene expression from scRNA-seq data

For individual genes, we considered the normalized gene expression across the cell populations. For each selected gene, we compared its expression within the 35 d cells with the one within the 70 d cells by computing the logarithmic fold change (log2FC). We performed this analysis for the neuron specific genes (Figure 1F) and neuronal subtypes including glutamatergic neurons, GABAergic neurons, and dopaminergic neurons (Figure 2B-D) where negative values indicate that a gene is less expressed at day 35 than at day 70 and positive numbers the opposite. p-value are based on Z-test with Bonferroni correction and significance levels correspond to * = p-value < 0.05, ** = p-value < 0.01, *** = p-value < 0.001, **** = p-value < 0.0001. Error bars represent SEM based on the individual sample average and error propagation.

### TEM Morphology

63 day old hMO specimens were immersion-fixed in a solution of 2 % PFA and 2.5 % glutaraldehyde in 0.1 M sodium cacodylate buffer (pH 7.4, Electron Microscopy Sciences, Hatfield, PA) for 3 h, rinsed several times in cacodylate buffer and further post-fixed in 2 % glutaraldehyde in 0.1 M sodium cacodylate buffer for 2 h at room temperature on a gentle rotator; fixative was allowed to infiltrate an additional 48 h at 4 °C. Specimens were rinsed several times in cacodylate buffer, post-fixed in 1.0 % osmium tetroxide for 1 h at room temperature and rinsed several times in cacodylate buffer. Samples were then dehydrated through a graded series of ethanols to 100 % and dehydrated briefly in 100 % propylene oxide. Tissue was then allowed to pre-infiltrate 2 h in a 2:1 mix of propylene oxide and Eponate resin (Ted Pella, Redding, CA), then transferred into a 1:1 mix of propylene oxide and Eponate resin and allowed to infiltrate overnight on a gentle rotator. The following day, specimens were transferred into a 2:1 mix of Eponate resin and propylene oxide for a minimum of 2 h, allowed to infiltrate in fresh 100 % Eponate resin for several hours, and embedded in fresh 100 % Eponate in flat molds; polymerization occurred within 24-48 h at 60 °C. Thin (70 nm) sections were cut using a Leica EM UC7 ultramicrotome, collected onto formvar-coated grids, stained with uranyl acetate and Reynold’s lead citrate and examined in a JEOL JEM 1011 transmission electron microscope at 80 kV. Images were collected using an AMT digital imaging system with proprietary image capture software (Advanced Microscopy Techniques, Danvers, MA).

### Microelectrode array

The Maestro microelectrode array (MEA, Axion BioSystems) platform was used to record spontaneous activity of the hMOs. A 48-well MEA plate containing a 16-electrode array per well was precoated with 0.1 mg/ml poly-D-lysine hydrobromide (Sigma-Aldrich). 60-70 days old organoids of two different passages were briefly treated for 5 min with 1X TrypLE Select Enzyme, resuspend in 10 µg/ml laminin (Sigma-Aldrich) and placed as a droplet onto the array. After 1 h incubation, neuronal maturation media was added and cells were cultured for 1-2 weeks. Spontaneous activity was recorded at a sampling rate of 12.5 kHz for 5 min at 37 °C over several days. Axion Integrated Studio (AxIS 2.1) was used to assay creation and analysis. A Butterworth band pass filter with 200-3000 Hz cutoff frequency and a threshold of 6x SD were set to minimise both false-positives and missed detections. The spike raster plots were analysed using the Neural Metric Tool (Axion BioSystems). Electrodes with an average of ≥5 spikes/min were defined as active, for the pharmacological treatment 24 electrodes were analysed. The organoids were consecutively treated with Gabazine, D-AP-5, NBQX (Cayman Chemical, end concentration: 50 mM each), and Quinpirole (Sigma Aldrich, end concentration: 5µM). To block all neuronal activity and thus verify spontaneous spiking activity of the cells, tetrodotoxin (TTX, Cayman Chemical, 1 µM) was applied at the end. The spike count files generated from the recordings were used to calculate the number of spikes/active electrode/minute. Further details regarding the MEA system were previously described (Bardy et al., 2015).

### Statistical analyses

If not stated otherwise, experiments were performed with three independently generated organoid cultures from three different cell lines (n=9). Gaussian distribution was evaluated by performing D’Agostino & Pearson omnibus normality test. In case the data were normally distributed, Grubbs’ test was performed to detect significant outliers. Unpaired t-test with Welch’s correction or nonparametric Kolmogorov-Smirnov test were performed to evaluate statistical significance. Data are presented as mean ±SEM. The statistical analyses of scRNA-seq data are described in the corresponding sections.

## Results

### Characterisation of the neuronal differentiation dynamics in midbrain-specific organoids

Previously, we demonstrated that human iPSC-derived midbrain floor plate neural progenitor cells (mfNPCs) can give rise to 3D human organoids that contain high amounts of dopaminergic neurons (Smits et al., 2019). To have a better insight into the dynamics of the neuronal differentiation, we evaluated TUJ1 staining, as a marker for neuronal differentiation, at two time-points during the differentiation of hMOs (Figure 1A). An in-house developed image analysis algorithm was used to segment Hoechst positive nuclei and TUJ1 positive neurons to create specific nuclear and neuronal masks. These masks contain all positive pixel counts for Hoechst and TUJ1, respectively.

The TUJ1 signal normalised to the Hoechst signal significantly increased after 70 days compared to 35 days, demonstrating a progressive differentiation into post-mitotic neurons. Whereas, the nuclear marker signal was significantly decreased at 70 days compared to 35 days, which might indicate selection in the cell population, as reported by Suzanne and Steller (2013) (Figure 1B). Along with these findings, we observed that the size of the organoids significantly increased during the course of the differentiation. This suggests that the increased TUJ1 volume and organoid size are due to the increased tissue complexity (e.g. neuronal arborisation) within the hMO (Figure 1B).

To further characterise the neuronal differentiation dynamics at the gene expression level, we performed scRNA-seq on samples from the two time-points mentioned above. The experiments were conducted using the Drop-Seq technique (Macosko et al., 2015), and the standard bioinformatics processing of the data resulted in two sample specific digital expression (DEM) matrices, which were further normalised and merged (see Methods section).

To visualise the so-obtained high-dimensional single-cell data, we performed dimensionality reduction of the DEM by t-distributed stochastic neighbourhood embedding (t-SNE) (van der Maarten and Hinton 2008) where each dot corresponds to a cell (Figure 1C). This distribution shows that cells originating from organoids at 35 days and 70 days only partially cluster together. This underlines that there are remarkable differences in the overall gene expression between the samples at days 35 and 70. The distribution of the cumulative gene expression (right panel in Figure 1C) shows that the neuronal gene expression is increased in a large fraction of the cells suggesting that this fraction of cells may not represent mature neurons.

To determine the differences in neuron-specific cumulative gene expression over time, we plotted the distributions of cells across the cumulative gene expression scores (Figure 1D). Here, we used the scores obtained from the lists of precursor cell-specific genes (indicated as “stemness genes”) and those of neurons. While the distributions of cells across the cumulative gene expression for precursors is very similar between day 35 and 70 (upper panel), we observed a significant difference between the distributions of cells across the neuronal cumulative gene expression (lower panel). Cells at 35 days tend to express the neuron-specific genes significantly more than cells at 70 days.

To further investigate how the differentiation of precursor cells into neurons evolves over time, we computed the gene-gene correlation for the genes of the neuron-specific list and of the stemness-specific list, altogether. Comparing these two lists, we found that at 35 days there are low values of correlation between genes exclusively specific for neurons or stem cells and also between neuron-and stemness-specific genes (Figure 1E, left heatmap). Very few of the correlation values are significantly different from zero and were substituted by zeros in the upper triangular matrix (Figure 1E, left heatmap). While correlations between stemness genes and neuron-stemness correlations remain similar at day 70 to day 35, correlations between neuron-specific genes increased considerably at day 70. This significant increase of neuron-specific gene correlations indicates a higher commitment of the cells towards the neuronal fate at day 70 compared to day 35 and supports the finding of a progressive maturation of post-mitotic neurons (Figure 1E).

Next, we wanted to elucidate which individual genes contribute to the differences between the gene expression of cells at day 35 and cells at day 70. For this purpose, we performed an analysis of DEGs across the whole transcriptome. From the 24,976 distinct transcripts measured, 1,311 were significantly differentially expressed between day 35 and 70 (p-value < 0.01 after Bonferroni correction, which represents approximately 5 % of all genes expressed (see Supplementary Table 1). When intersecting the list of DEGs with the stemness-specific genes, we found that approximately 30 % of the stemness-specific genes are DEGs (see Table 4). Similarly, 42 % (corresponding to 11 genes) of the neuron-specific genes are differentially expressed (Table 3 and 4). Hence, the percentages of DEGs within the neuron-specific and stemness-specific lists are remarkably higher than the percentage of DEGs across the whole transcriptome. These notable changes further indicates the induction of the neuronal differentiation and progressive maturation.

**Table 4:** List of differentially expressed genes.

| Stemness | Neuronal | Dopaminergic | Glutamatergic | GABAergic | Serotonergic |
| --- | --- | --- | --- | --- | --- |
| ASCL1 | CALB2 | APP | GRIA2 | GAD2 | HTRA1 |
| FOXA2 | CELF4 | EN1 | GRIN2A |  | SLC18A24 |
| LMO3 | CLSTN2 | FOXA2 | GRIN2B |  |  |
| SOX2 | DLX1 | GPC2 | SLC17A6 |  |  |
|  | DPYSL5 | NTM | SLC1A2 |  |  |
|  | EBF3 | PCDH10 | SLC1A3 |  |  |
|  | FOSL2 | PDZRN4 |  |  |  |
|  | SLC17A6 | TH |  |  |  |
|  | SST |  |  |  |  |
|  | STMN2 |  |  |  |  |
|  | VSTM2L |  |  |  |  |

Next, we focused on individual neuron-specific genes and compared their expression over time. For each gene we computed the log2 fold-change between the average expression at day 35 and day 70 (Figure 1F). Consistent with the findings shown in Figure 1D, the majority of genes were significantly higher expressed at day 35 than at day 70 based on Z-test analysis corrected for multiple hypotheses testing. Interestingly, among the genes expressed at day 35 we found genes that are involved in neurogenesis (EBF3 (Garcia-Dominguez et al., 2003)), neuronal migration and differentiation (L1CAM (Patzke et al., 2016)), whereas genes expressed at day 70 reflect more specific subpopulations of neurons, like GABAergic neurons (DLX1, CALB2 (Al-Jaberi et al., 2015)) in agreement with the separate DEG analysis across the whole transcriptome (see Table 4).

### Midbrain-specific organoids consist of different neuronal subtypes

From previous studies we know that hMOs are rich in dopaminergic neurons (Jo et al., 2016; Qian et al., 2016; Monzel et al., 2017; Smits et al., 2019; Kim et al., 2019). We wanted to further explore which other neuronal subtypes develop besides midbrain dopaminergic neurons within the hMOs.

Therefore, we investigated the expression of genes typical for dopaminergic, glutamatergic, GABAergic, and serotonergic neurons by analysing the scRNA-seq data. We plotted the distributions of cells across the cumulative gene expression scores, which were obtained from the lists of genes specific of a neuronal subtype (Figure 2A). While the cell distribution over cumulative expression score for GABAergic neurons was very similar between the samples at 35 days and 70 days (Figure 2A, third panel), we detected statistically significant differences between the distributions of cells over scores for the other three types of neurons. The expression of the selected genes for the glutamatergic and dopaminergic neurons was increased at day 35 compared to day 70, which is consistent with the observations for the neuron specific score (Figure 1D). Interestingly, we discovered that the vast majority, approximately more than 700 of the 1000 cells, lacks completely expression of genes specific for serotonergic neurons (Figure 2C, fourth panel). Thus, we disregarded this neuronal subtype in the subsequent analyses and focused next on individual genes specific of dopaminergic, glutamatergic, and GABAergic neurons, by computing the log2 fold-change between the average gene expression at day 35 and at day 70 (Figure 2B-D). In each of the three lists, the majority of the genes for which a statistically significant difference is present are actually more expressed at 35 days than at 70 days, consistently with the findings of Figure 2A.

When intersecting the list of 1,311 DEGs across the whole transcriptome with the lists of dopaminergic, glutamatergic and GABAergic neurons, we found that 42 %, 34 %, and 17 % of the genes were DEGs within these lists, respectively (see Supplementary Table 1 and Table 4). Again, all of these percentages are considerably higher than the 5 % of DEGs across the whole transcriptome indicating that neuronal subtypes specific genes, in particular dopaminergic and glutamatergic, are highly represented within DEGs during hMO development.

Lastly, to verify the presence of the addressed neuronal subtypes we conducted an immunohistochemistry staining for the respective neurotransmitters. This allowed us to robustly detect dopaminergic, glutamatergic and GABAergic neurons as well as even a few serotonergic neurons within hMOs (Figure 2E).

### Midbrain-specific organoids express synaptic proteins

After identifying the presence of neurons and even specific neuronal subtypes on transcriptome expression levels by means of neurotransmitter staining and scRNA-seq, we investigated the actual interaction among the neuronal cells within the hMOs. We previously showed that hMOs synthesise and release the neurotransmitter dopamine (Smits et al., 2019). This already suggests the establishment of a functional neuronal network. The basic requirement for neuronal network formation is the development of synapses. Hence, we evaluated the presence of synaptic connections using the presynaptic marker SYNAPTOPHYSIN and the postsynaptic marker PSD95 in organoid sections after 70 days of culture (Figure 3A). Both proteins were detectable in a puncta-like organisation, which is expected for synapses. With a subsequent 3D surface reconstruction, we observed that the signals for SYNAPTOPHYSIN and PSD95 were localised in close proximity, forming pre- and postsynaptic puncta (Figure 3B). To further investigate whether actual functional synaptic connections were formed in the hMOs, we used a transmission electron microscopy (TEM) approach (Figure 3C). EM micrographs show excitatory synapses characterised by electron dense post-synaptic density proteins (full arrow) and pre-synaptic synapse (asterisks) loaded with synaptic vesicles.

**Figure 3:**
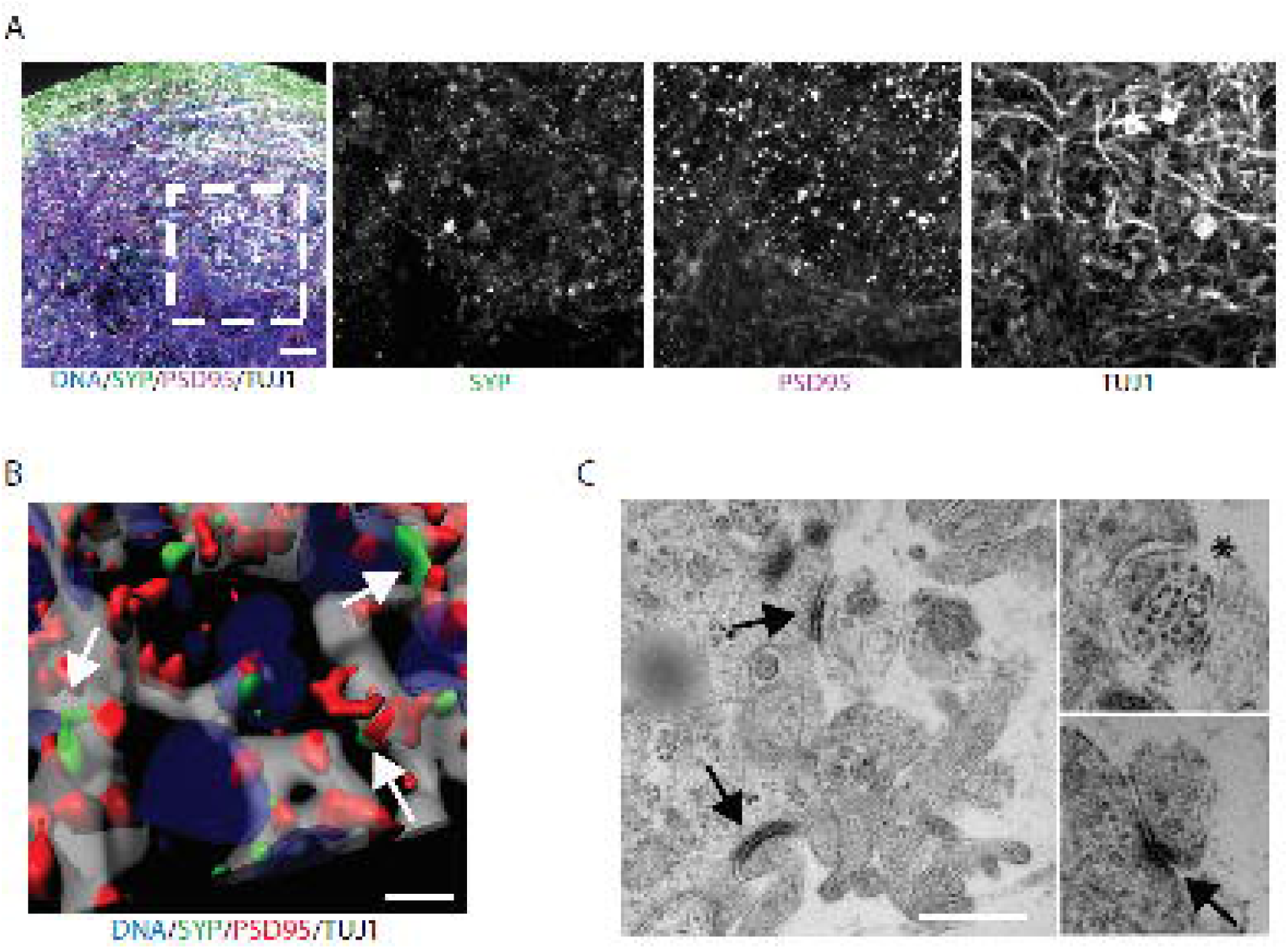
Midbrain-specific organoids express synaptic proteins. (A) Immunostaining of pre-and the postsynaptic markers at day 70. Dashed lines indicate the region of magnification. Scale bar is 50 µm. (B) 3D surface reconstructions of confocal z-stacks of an organoid at day 70 of differentiation. Scale bar is 10 µm. (C) Representative electron micrographs of synaptic terminals from 63 d organoids. Scale bar is 500 nm.

### Midbrain-specific organoids develop GABAergic, glutamatergic and dopaminergic electrophysiological activity

Non-invasive multielectrode array (MEA) measurements can give insights into physiological properties, like the generation of spontaneous neuronal activity of *in vitro* cultured, self-organised networks (Luhmann et al., 2016). As the assessment of neuronal activity is important to evaluate the functional maturation, we tested the spontaneous electrophysiological activity of hMOs by MEA measurements (Odawara et al., 2016). We measured extracellular field potentials, which are generated by action potentials. At days 50-60 of differentiation, hMOs were seeded in 48-well tissue culture MEA plates on a grid of 16 electrodes (Figure 4A and B). After 10-20 days of culturing, we recorded spontaneous activity, on several electrodes, over several days, in the form of mono- and biphasic spikes (Figure 4Aii). To investigate which neuronal subtypes were functionally active in the hMOs, we applied specific drugs following a previously reported experimental design (Illes et al., 2014). We recorded spiking patterns from 24 active electrodes: in Figure 4C and D representative recordings of one electrode are displayed. After treating the organoids with gabazine, a GABA_A_ receptor antagonist, we detected an increase of spontaneous spiking (22.5 % increase, Figure 4Dii). Following the gabazine-induced disinhibition, we applied the AMPA/Kainate-receptor antagonist NBQX and the NMDA-receptor antagonist D-AP-5. The inhibition of the excitatory neurons resulted in a 28.1 % decrease of spontaneous activity (Figure 4Diii). After the inhibition of GABAergic and glutamatergic neurons in the hMOs, we added the D2/D3 receptor agonist quinpirole (Figure 4C and Div), which resulted in a 47.8 % decrease of neuronal activity. Confirming the findings displayed in Figure 2, we conclude from these experiments that hMOs contain functional GABAergic, glutamatergic and dopaminergic neurons.

**Figure 4:**
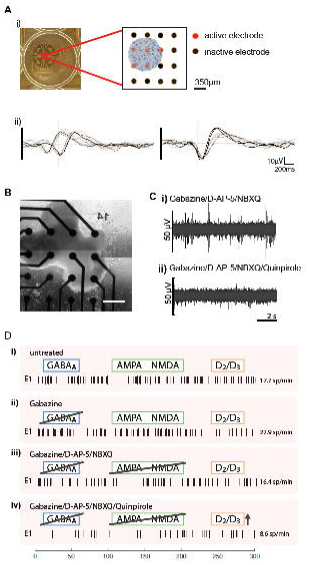
Electrophysiological activity in midbrain-specific organoids. (A) Representative scheme of positioned midbrain organoid on a 16-electrode array in a 48-well tissue culture plate (i). Examples of mono-and biphasic spikes detected by individual electrodes of a multielectrode array (MEA) system (ii). (B) Representative image of midbrain organoid positioned on a 16-electrode array in a 48-well tissue culture plate. Scale bar is 350 µm. (C-D) Evaluation of the spontaneous activity by addressing inhibitory (blue) and excitatory (green) neurotransmitter receptors using multielectrode array (MEA) system. (C) Representative raw data traces show the effect of Quinpirole in absence of inhibitory and excitatory synaptic communication. (D) Representative spik raster plots demonstrate effects of applied compounds.

## Discussion

The *in vitro* human brain organoid technology has become a valuable tool allowing advances in the field of basic research as well as in translational applications (Fatehullah et al., 2016). Organoids specifically modelling the human midbrain hold great promise for studying human development and for modelling Parkinson’s disease (PD) (Jo et al., 2016; Monzel et al., 2017; Kim et al., 2019; Smits et al., 2019). In contrast to 2D monolayer cultures, hMOs can recapitulate complex interactions of midbrain dopaminergic neurons with other cell types of the central nervous system (CNS) in a 3D environment. However, human midbrain brain organoid research has so far focused mainly on dopaminergic neurons. In a detailed study of Borroto-Escuela et al. (2018) it has been described that released dopamine can diffuse into synaptic regions of glutamate and GABA synapses and directly affect other striatal cell types possessing dopamine receptors. Furthermore, *substantia nigra* dopaminergic neurons are directly controlled by GABAergic input (Tepper and Lee, 2007). Evidences from these studies suggest that the presence of other neuronal subtypes is important to model multifactorial disease like PD. In our study, we have demonstrated that the derivation of hMOs leads to functional neuronal networks, containing different neuronal subtypes of the human midbrain. Single-cell transcriptomic data from hMOs demonstrated that there is an increased expression of neuronal-specific genes in 35 days compared to 70 days old hMOs. On the other hand, the gene-gene correlations between only neuron-specific genes increased considerably at day 70, suggesting an increased commitment of cells towards the neuronal cell fate during the course of the organoid development. This further supports the finding of a progressive maturation of post-mitotic neurons (Figure 1D and E). The identification of these neuron-specific genes revealed that the genes upregulated at the earlier time point are relevant in the process of neurogenesis and neuronal migration and differentiation (EBF3 (Garcia-Dominguez et al., 2003), L1CAM (Patzke et al., 2016)). Whereas the upregulated genes at the later time point have been for instance implicated in subpopulations like GABAergic neurons (DLX1, CALB2 (Al-Jaberi et al., 2015)). (Figure 1F). This indicates a higher commitment of the cells toward their intended fate and a progressive maturation of the post-mitotic neurons within the hMOs. Furthermore, single-cell analysis of the hMOs also proved the presence of specific neuronal subtypes, like dopaminergic, glutamatergic and GABAergic neurons. While the scRNA-seq data were not fully conclusive concerning serotonergic neurons, a staining approach allowed to detect at least some of these neurons (Figure 2). Supporting the findings of currently published midbrain-specific organoid models (Jo et al., 2016; Qian et al., 2016; Monzel et al., 2017; Smits et al., 2019), we detected a significant upregulation of tyrosine hydroxylase (TH) within the cell population of 70 days old hMOs compared to 35 days old hMOs. The activity of neurons and their different receptors can be analysed by the specific response to chemical compounds. It has been shown that quinpirole, a specific D2/D3 receptor agonist, suppresses the firing in hMOs (Jo et al., 2016; Monzel et al., 2017). In addition to the previously reported analyses in hMOs, we blocked inhibitory and excitatory communication, to further isolate and attribute the recorded signals to neuronal subtypes. Gabazine induces a disinhibition of GABAergic neurons, whereas NMDA-receptor and AMPA/Kainate-receptor antagonists inhibit glutamatergic excitatory communication (Illes et al., 2014). Together with the characteristic hallmarks of synapse formation (Figure 3A-C) and the previous findings of dopamine release (Smits et al., 2019), these data confirm the presence of functional dopamine receptors in dopaminergic neurons as well as functional GABAergic and glutamatergic neurons within hMOs. As neurons do not exist in isolation in the CNS but form functional networks with other neurons and non-neuronal cells, it is important to expand our research of neurodegenerative diseases using 3D models that are able to recapitulate cell autonomous as well as non-cell autonomous aspects. Utilising 3D cell culture models that comprise a variety of neuronal subtypes could lead to new insights into the selective vulnerabilities, which are observed in neurodegeneration. Indeed, evidence suggests that specific regulation of the excitability of dopaminergic neurons by other neuronal subtypes in the midbrain might explain their selective vulnerability in PD (Korotkova et al., 2004). This underlines the importance and the enormous potential for future disease modelling of the here described hMO model, as it contains functionally connected heterogeneous neuronal cell populations.

## Acknowledgments

We would like to thank Dr. Sebastian Illes (University of Gothenburg, Sweden) for his help with the experimental MEA design and Diane Capen (Massachusetts General Hospital, Boston, USA) for her EM work. We thank Dr. Jared Sterneckert (Technical University of Dresden, Germany) and Dr. Bill Skarnes (The Wellcome Trust Sanger Institute, Cambridge, UK) for human iPSC lines. Furthermore, we would like to thank Yohan Jarosz from the LCSB Bioinformatics Core Group for his support in data management. This project was supported by the LCSB pluripotent stem cell core facility. The JCS lab is supported by the Fonds National de la Recherché (FNR) (CORE, C13/BM/5791363 and Proof-of-Concept program PoC15/11180855 & PoC16/11559169). This is an EU Joint Programme-Neurodegenerative Disease Research (JPND) project (INTER/JPND/14/02; INTER/JPND/15/11092422). Further support comes from the SysMedPD project which has received funding from the European Union’s Horizon 2020 research and innovation programme under grant agreement No 668738. LMS is supported by fellowships from the FNR (AFR, Aides à la Formation-Recherche). Electron microscopy was performed in the Microscopy Core of the Center for Systems Biology/Program in Membrane Biology, which is partially supported by an Inflammatory Bowel Disease Grant DK043351 and a Boston Area Diabetes and Endocrinology Research Center (BADERC) Award DK057521. SM is supported by the University of Luxembourg and the National Research Fund through the CriTiCS DTU FNR PRIDE/10907093/CRITICS. KG and AS were supported by the Luxembourg National Research Fund (FNR) through the C14/BM/7975668/CaSCAD project and AS additionally by the National Biomedical Computation Resource (NBCR) through the NIH P41 GM103426 grant from the National Institutes of Health.

## Author Contributions

LMS designed and performed cell culture and imaging experiments, prepared the figures and wrote the original draft. KG performed the scRNA-seq experiments and related bioinformatics approaches. SM performed the computational analysis of the single-cell RNA-Seq data, edited the manuscript and contributed to the figures. PMAA contributed to the development of 3D image analysis. RK supervised image analysis design. AS supervised the design and implementation of the single-cell experiments and associated computational data analysis. SB initiated the project, supervised it and edited the manuscript. JCS conceived and supervised the project, designed the experiments and edited the manuscript.

## Data Availability

The data that support the findings of this study are public available at this doi: www.doi.org/10.17881/lcsb.20190326.01.

