## Supplementary Information for "Single-cell transcriptomics reveals multiple neuronal cell types in human midbrain-specific organoids"

**Corresponding author**

Prof. Dr. Jens C. Schwamborn

**Supplementary Table**

**Supplementary Table 1:** **Analysis of DEGs across the whole transcriptome**. 1,311 genes from 24,976 distinct transcripts measured, were significantly differentially expressed between day 35 and 70 (p-value < 0.01 after Bonferroni correction, representing approximately 5 % of all genes expressed).

This table is uploaded as a separate excel sheet.
